# Adaptive proximity to criticality underlies amplification of ultra-slow fluctuations during free recall

**DOI:** 10.1101/2023.02.24.529043

**Authors:** D. Yellin, N. Siegel, R. Malach, O. Shriki

## Abstract

Ultra-slow fluctuations are a hallmark of spontaneous cortical activity. We examine the hypothesis that these unique dynamics arise from recurrent neuronal networks operating near a phase transition, a state characterized by ‘critical slowing down’. A further prediction of such dynamics is that a small modulation towards the critical transition should lead to specific amplification of slow fluctuations. Here, we relate this phenomenon to experimental findings using a simulation of a simple random recurrent network. Importantly, the model aligns with direct intracranial electroencephalography recordings from human visual cortex during both rest and visual free-recall, specifically replicating the observed enhancement of slow fluctuations during free recall. These simulations illuminate a simple and powerful mechanism underlying slow spontaneous fluctuations, while enabling the rapid transition between different spontaneous states. They propose that modulation towards criticality might be a universal strategy employed by cortical networks to engage in a spontaneous generative mode.

## Introduction

Ultra-slow spontaneous activity fluctuations, also known as “resting-state” activity (Raichle, 2011), have been proposed to drive free and creative behaviors in the human brain (Broday-Dvir & Malach, 2021; Norman & Malach, 2022). The power spectrum of these fluctuations has been characterized as obeying a power-law profile (He, 2014; Miller et al., 2009; Nir et al., 2008) and they emerge in well-defined networks across the cortex, uncovering unique patterns of functional connectivity (Buzsáki, 2006; Fox & Raichle, 2007; Nir et al., 2006). These spontaneous fluctuations have also been linked to ramping anticipatory signals, as well as other spontaneous behavioral manifestations, such as pupil diameter fluctuations (Yellin et al., 2015) and eye-drifts (Ramot et al., 2011). We previously proposed (Moutard et al., 2015; Norman et al., 2017) and recently demonstrated, a direct link between the spontaneous activity fluctuations and free and creative verbal behaviors (Broday-Dvir & Malach, 2021; Malach, 2024). However, it is still not clear how the characteristic power spectrum of the spontaneous fluctuations - dominated by ultra-slow frequencies - emerges out of the typically fast neuronal elements. We have previously suggested that these slow dynamics emerge as a result of recurrent neuronal connections (Moutard et al, 2015).

### Near-critical dynamics as a potential mechanism

Here, we extend this notion and examine the hypothesis that the cortex operates in the vicinity of a critical transition point (Beggs, 2022; Beggs & Plenz, 2003; Kinouchi & Copelli, 2006; Shew et al., 2009). In such a system, the slow resting state fluctuations can be explained in the context of a fundamental property of near-critical recurrent systems - the slowing down of stochastic fluctuations as these systems approach criticality. Termed “critical slowing down” (Lee et al., 2014; Meisel et al., 2015; Scheffer et al., 2009), the phenomenon describes the universal tendency of systems near a transition point in their dynamics to exhibit slower time scales and take longer time to relax to a steady state equilibrium following perturbations. Hence, the slowing down of spontaneous fluctuations near criticality may explain their unique power spectra as observed in the resting state networks. Indeed, some prior modeling work has tentatively proposed such a link (Chaudhuri et al., 2018; Deco & Jirsa, 2012).

Recent experimental and theoretical evidence supports the idea that the healthy cortex operates near a critical point, exhibiting slow time scales and scale-free dynamics (Arviv et al., 2015; Deco & Jirsa, 2012; Haimovici et al., 2013; Linkenkaer-Hansen et al., 2001; Shew & Plenz, 2013; Shriki et al., 2013; Yu et al., 2013). Scale-free spatiotemporal cascades of neuronal activity, termed neuronal avalanches, have been found in *in-vitro* and *in-vivo* animal studies as well as in large-scale human fMRI, EEG and MEG recordings (Arviv et al., 2015; Lombardi et al., 2021; Palva et al., 2013; Tagliazucchi et al., 2012). These avalanches can be well described by the framework of critical branching processes (Harris, 2012) and they can coexist with oscillations (Lombardi et al., 2023). Other studies indicated edge-of-chaos criticality (Bertschinger & Natschläger, 2004; Legenstein & Maass, 2007; Toker et al., 2022). Another type of critical transitions discussed in the literature is pattern formation critically, which refers to a transition into self-organized spatial patterns of activity, such as stationary bumps or traveling waves (Bressloff & Cowan, 2003; Chossat & Faugeras, 2009; Pinto & Ermentrout, 2001; Shriki & Yellin, 2016).

Importantly, near-critical dynamics optimize information processing (Shew et al., 2009; Shriki & Yellin, 2016) and deviations from critical dynamics are associated with disorders in information processing (Arviv et al., 2016; Dotan & Shriki, 2021; Fekete et al., 2018, 2021). We have previously demonstrated the phenomenon of critical slowing down in a recurrent network model of early visual processing, where the effective time scales grow longer as the network approaches a critical point during its learning process (Shriki & Yellin, 2016). For a recent review of critical brain dynamics, see (Beggs, 2022; O’Byrne & Jerbi, 2022).

A straightforward prediction of the near-critical slowing down dynamics is that bringing the system closer to criticality should result in biasing the power spectra of the spontaneous fluctuations towards even slower frequencies. Indeed, recent experimental findings derived from direct recordings of brain activity in patients are compatible with this notion. Thus, in a study exploring brain mechanisms of visual recall, Norman et al. (2017) found that resting state fluctuations displayed an increase in baseline neuronal firing rate and power of the slow spontaneous fluctuations during free recall. This local baseline shift - i.e., increase in average firing rate - was predicted to bring the targeted neurons closer to the decision threshold and enhance the probability of free recall during such periods.

It is important to emphasize that this amplification of the fluctuation amplitude can serve a significant cognitive and functional role. Free behavior relies on the ability to rapidly and flexibly switch between active and rest conditions. For instance, musicians can easily switch into free improvisation mode from a state of rest. If the slow spontaneous fluctuations indeed underlie free behaviors, a mechanism must be present in the brain to regulate these fluctuations. In the study of Norman et al (2017), exploring this idea in the domain of free visual imagery and recall, it was found that resting state fluctuations displayed an increase in baseline and power of the slow spontaneous fluctuations specifically when subjects switched from rest to an active recall state. Here, we propose that this local dynamical shift, which increases the probability of targeted neurons to cross the decision threshold, can be elegantly explained by the critical slowing down phenomenon.

Specifically, we propose that the related amplification of the spontaneous fluctuations may be due to bringing the network closer to its critical point, leading to critical slowing down. This provides a mechanism for the efficient implementation of a state-change from rest to active, yet, spontaneous behaviors. We further speculate that the proximity to criticality is controlled through neuromodulation, allowing dynamic regulation of cortical states

### Network model and tuning of proximity to criticality

To explore the suggested mechanism, we simulated a random recurrent network of rate-based units at varying proximities to criticality. These rate models can be rigorously derived from large-scale conductance-based networks that model cortical dynamics (Shriki et al., 2003). Our network architecture is adapted from the one proposed by Chaudhuri et al. (2018) to simulate intra-cranial EEG (iEEG) dynamics. Proximity to criticality is controlled by tuning relevant biophysically meaningful parameters. The precise tuning is determined by analyzing the recurrent interaction matrix using considerations from random matrix theory assuming a large enough network size, typically in the order of a few hundred neurons (Ganguli et al., 2008; Rajan & Abbott, 2006; Vázquez-Rodríguez et al., 2017). Near-criticality is required to reproduce the observed scale-free power spectrum of iEEG recordings (Chaudhuri et al., 2018). Setting the model closer to its critical point may account for the unique dynamics observed during free recall (Norman et al., 2017).

To provide further insight, Fig. 1 schematically illustrates critical slowing down (CSD) in a random recurrent neural network and in a one-dimensional non-linear system near its critical point. Panel 1a shows the expected activity in a random recurrent network at varying proximities to its critical point, as set by the network gain parameter, *G*. The critical point is defined when *G* = 1 (see Methods for further mathematical substantiation). For a transient stimulus, activity will dissipate when the network gain is set to *G* < 1 (blue arrow), diverge to the maximal neuronal rate (in black dashed line) when *G* > 1 (green arrow), and tend to persist at enhanced slow fluctuation frequencies when *G* ≲ 1, namely slightly below the critical point (yellow arrow). For a continuous signal, the steady-state power spectrum indicates an increase in the amplitude of slow activity only for *G* ≲ 1.

**Figure 1.**
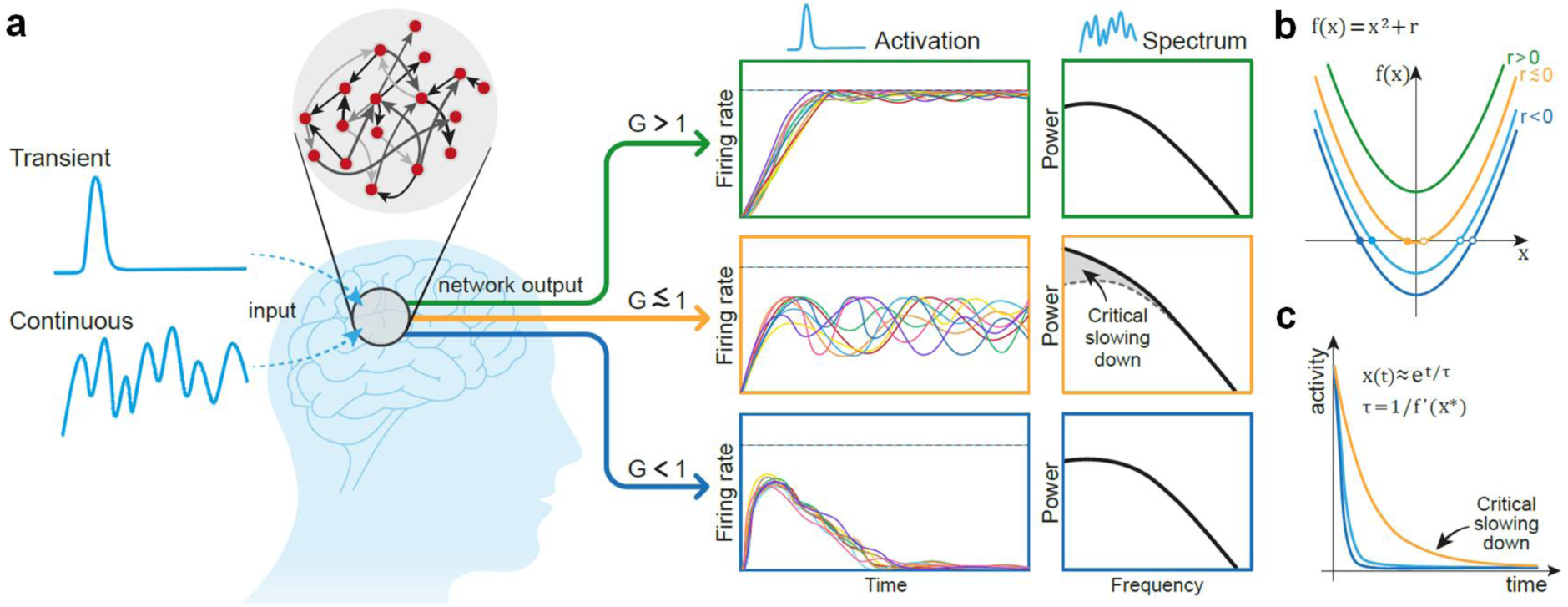
The impact of gain change on the dynamics of a recurrent system. The figure provides an illustrative and simplifying explanation of the critical slowing down (CSD) phenomenon in the dynamics of a recurrent system. CSD takes place as the system crosses between its sub-critical domain (G < 1, here depicted in blue) and its super-critical domain (G > 1, here in green). In this near-critical state (G ≲ 1, here depicted in yellow), when the system’s gain (also termed ‘control’) parameter is increased to the vicinity of (but still below) the critical point, it displays cardinal slowing down and amplification of its dynamics. Panel 1a delineates the expected activity in a random recurrent network at varying proximity to its critical point. For a transient stimulus, activity in the network will (a) dissipate in the sub-critical domain (G < 1, in the bottom sub-panel) (b) diverge to the maximal neuronal rate (black dashed line) in the super-critical domain (G > 1, top sub-panel) and (c) tend to become persistent as the gain approaches G ≲ 1 (middle sub-panel). For a continuous signal, the power spectrum will indicate rise in the amplitude of slow-wave activity only at the domain of G ≲ 1. Panels 1b and 1c provide further illustration of the CSD phenomenon exploring the dynamics of a simple one-dimensional non-linear system, driven by the quadratic equation of f(*x*) = *x*^2^ + *r*. Panel 1b depicts the system and its solutions as the value of *r*, the control parameter is modified between super-critical (r > 0, in green), sub-critical (r < 0, blue) and near-critical (yellow, r ≲ 0) states. Panel 1c shows how the system dynamics slows down as r approaches 0. The curve corresponding to r > 0 diverges exponentially (not shown). See a more detailed discussion in the text.

Panels 1b & 1c further illustrate the CSD phenomenon in a one-dimensional system. The dynamics are characterized by a simple quadratic equation: f(*x*) = *x*^2^ + *r*, where *x* is a time-dependent variable satisfying 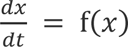, and *r* is the system’s control parameter. At *r* = 0 the system has a critical (bifurcation) point. For *r* > 0 it is super-critical, since at this domain there are no fixed points and *x* always diverges to infinity, whereas at *r* < 0 it is sub-critical and two fixed points arise at 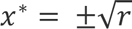 (panel 1b; filled circles correspond to stable fixed points and hollow circles to unstable fixed points). Close to a given fixed point, the dynamics can be linearized and the deviation from the fixed point can be approximated by an exponential temporal profile, e^t/τ^, where *τ* = 1/f′(*x*^∗^) (note that for stable fixed-points the derivative is negative and the dynamics converge). The magnitude of the time constant, *τ*, that controls the dynamics approaches infinity as the system approaches its critical point (where f ^′^(*x*^∗^) = 0). Thus, the dynamics slow down as the system approaches the critical point (panel 1c). Furthermore, at the bifurcation, the linear term vanishes, and the dynamics become dominated by higher-order terms, leading to power-law scaling behavior. This transition to power-law dynamics is a hallmark of critical systems, reflecting the absence of a characteristic timescale and the emergence of scale-invariant fluctuations

In summary, we propose that the unique dynamics observed by Norman et al. (2017) may be accounted for by a closer proximity of the local cortical network to a critical point during free recall. This enhances the gain of the local network, making it more sensitive to noise and driving it closer to the decision bound. We propose a model of a randomly connected recurrent network subject to a white noise input for this putative mechanism. Our key hypothesis is that this amplification in the slow activity amplitude stems from the critical slowing down phenomenon, and can be obtained by a relatively small shift towards criticality. It is likely that any model that manifests such critical phase transition behavior and power law spectra would be equivalently sufficient. Our findings support the hypothesized slowing down modulation near-criticality, showing good fit with the empirical data. They support the hypothesis that the phenomenon of ultra-slow, scale-free resting state fluctuations and their modulation during intrinsically motivated tasks reflect a basic and ubiquitous behavior common to all suitably configured recurrent systems.

## Results

To model the iEEG signal, we simulated a sparse random recurrent network, realizing a firing rate model (see Methods). The summed activity of a sample of units represented the corresponding iEEG. The proximity to the critical transition point between stable and unstable dynamics can be evaluated using tools from random matrix theory, which rely on the assumption of a large network size. To satisfy the requirement for a large enough neuron population, we set the network size (N) to 240 units (see Methods and SI Fig. S1 for a detailed analysis). Specifically, the proximity to criticality was directly modulated through a control parameter, G, corresponding to the product of the network sparseness p, the neural gain, *`γ`*, and the mean synaptic strength *`μ`*_conn_, with the critical transition from subcritical to supercritical dynamics expected at G_crit_ = *`γ`*p*`μ`*_conn_ = 1 (Methods). Although the values of p, *`γ`*, and *`μ`*_conn_ that comprise G can vary to satisfy the criticality condition, for simplicity, we followed Chaudhuri et al. (2018), setting p = 0.2 and *`μ`*conn = 49.881 across our main simulations. The neural gain (*`γ`*) was adjusted to control the network’s proximity to the critical phase transition. The time constant (τ) in these simulations was set to 20 ms, which is close to well-known estimates for cortical neurons’ membrane time constant (Gao et al., 2020; Koch et al., 1996; Softky & Koch, 1993). The sampling fraction, α, for computing the iEEG signal, was set between 0.01 <= α <= 0.1 to better adjust the fit in mid-to high frequencies (> 1 Hz) where this was relevant (Chaudhuri et al., 2018). The mean and std of *I*^*ext*^, the noisy input, was ∼20 (±11.5) *pA*. To modulate the level of white noise, we used an additive white noise current, *I*^*add*^. Thus, only two parameters are actually tuned to optimize the slow frequency PSD fit – *`γ`* (influencing the value of G) and the additive white noise, *I*^*add*^ (see Methods and Fig 3).

We first examined whether our network model could reproduce the dynamical characteristics of the spontaneous resting state activity fluctuations as recorded in the human cortex, and found that it did when set close to its critical point. This is demonstrated in Fig. 2, which compares real iEEG recordings and simulated network dynamics. It has been previously shown (Manning et al., 2009; Mukamel et al., 2005; Nir et al., 2007) that fluctuations in the power of the LFP in the high frequency broadband (HFB) provide a reliable index of the fluctuations in the average firing rates of neurons within the recording site. Note that this HFB measure should be distinguished from the direct recordings of the LFP signal itself. Panel 2a shows high correlation levels (r = 0.56 Pearson) in spontaneous firing rate fluctuations based on HFB from two auditory cortex electrical contacts recorded across hemispheres during rest (obtained from Nir et al., 2008). The autocorrelation (black) and cross-correlation (orange) PSD of the HFB obtained from multiple auditory cortex sites across the two hemispheres during rest in Nir et al are shown in panel 2b.

**Figure 2.**
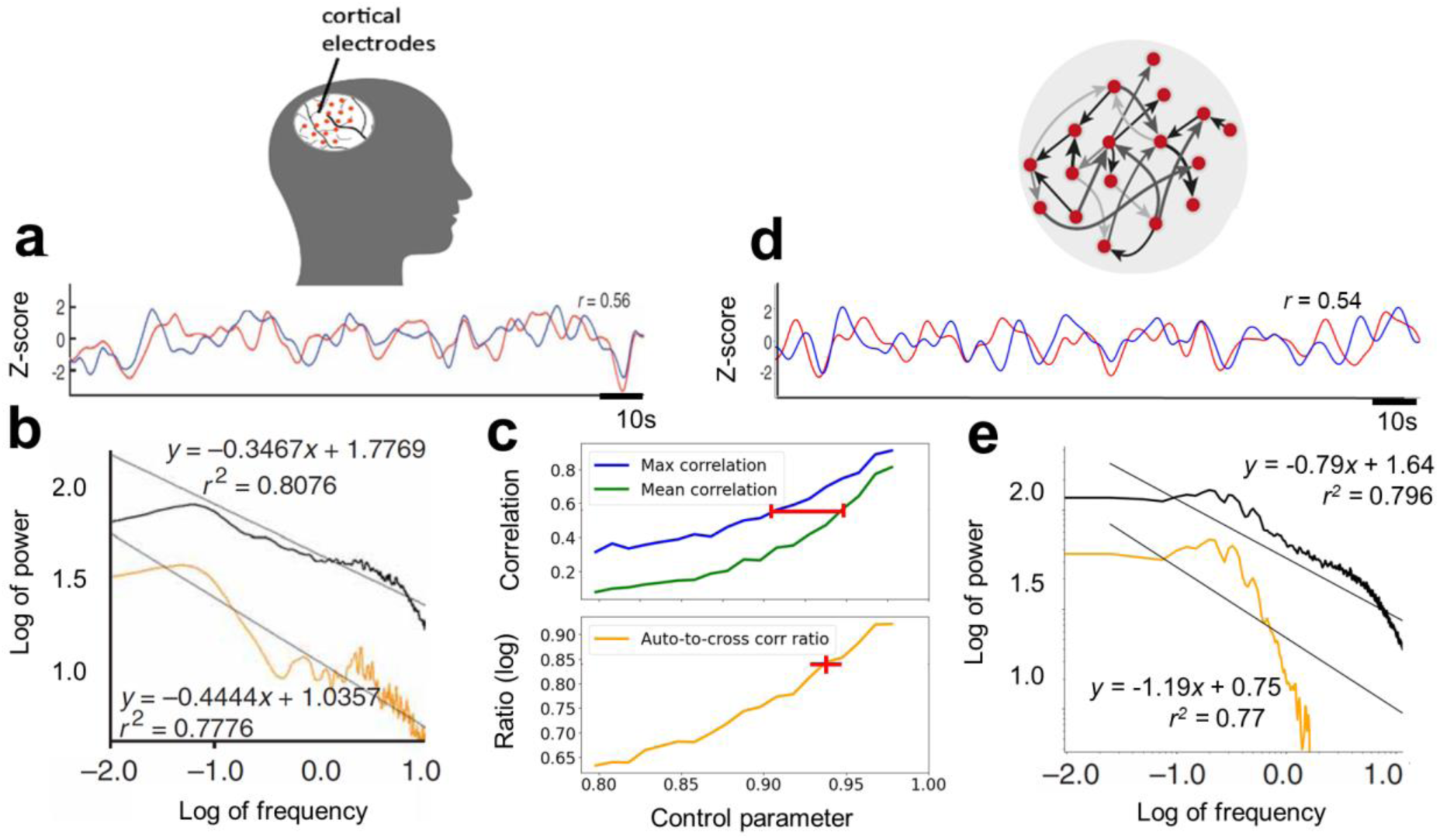
Ultra-slow spontaneous oscillations of neuronal activity in the brain and in a recurrent network simulation. Empirical iEEG results during wakeful-rest (adapted from Nir et al., 2008), presented here in panels 2a and 2b, are compared and contrasted with corresponding fitted simulation results in panels 2d and 2e, respectively, from an artificial neural network model with 240 units. Panel a presents the HFB signal from homological left and right auditory cortices. Panel 2b presents the autocorrelation (black line) and the cross-correlation (orange line) of the HFB PSD obtained across the two auditory cortex hemispheres. Panel 2c presents simulation analysis results placing computational bounds on expected values of the control parameter G, as it grows from the sub-critical to the critical range. The upper part of the panel shows the simulation results for maximum (blue line) and average (green line) pair correlation, placing a bound (red interval) on the G (at the range 0.91 – 0.96) for the simulated correlations to match empirical results in panel 2a. The lower part of panel 2c shows the log ratio of simulated auto-to cross-correlation mean power at low frequency (orange line) indicating best match to empirical results at G* ≈ 0.95∼ (red cross). Panel 2d demonstrates a corresponding signal from two highly correlated units in the artificial network at G*. Panel 2e depicts, for comparison, our simulation results of the activity fluctuations’ mean auto- and cross-correlation at G*. Note the similar location of the knee frequency in the autocorrelation spectra (at ∼1 Hz) of the empirical and simulated data (for further comparison see SI Fig. S2).

To investigate the influence of the proximity to criticality on the typical correlation levels between pairs of units in the network, we simulated multiple network realizations, gradually increasing the control parameter from the sub-critical (*G* = 0.8) to the near-critical (*G* = 0.99) range. The upper part of Panel 2c presents the simulation’s maximum and average pair correlation in blue and green, respectively. Results indicate that the network first reaches the correlation level observed by Nir et al only at *G* = 0.91 (blue curve), and that this becomes the mean correlation level at *G* = 0.96 (green curve), setting a bound on the operating range of *G* (see red interval).

Further “fine-tuning” of the actual value of the control parameter *G* can be estblished by computing the ratio of the power law exponents of these cross- and auto-correlation functions at low frequencies. The lower part of panel 2c shows the simulation results of this computation, in the range *G* = 0.8 to 0.99, over multiple realizations of the model network. Our findings show that the autocorrelation is less influenced by the growth of *G* relative to the cross-correlation, and thus, their exponent ratio changes rather steeply as a function of *G* (orange line), providing the approximation *G*\* ≈ 0.95 (see red cross) for the log ratio observed in the empirical findings (0.84). Panel 2d demonstrates correlated activity fluctuations of two simulated units in the recurrent network (see resemblance to Fig 2a), whereas panel 2e depicts the simulated auto- and cross-correlation PSD results for this estimated value *G*^∗^. For later reference, and given the inherent difficulty in determining an exact value for *G*^∗^, we define a typical resting-state range, *G*^∗^ = [0.94 − 0.95], within which *G*^∗^is expected to lie. Note that results for other values of τ were explored to optimize the fit of the “knee” bend in the PSD curve as presented in SI Fig. S2, showing that a τ value of 20 ms provides better fit than alternative settings at 2 or 200 ms. Note also that in the very near vicinity of the critical point (*G* ≥ 0.97) the slow mode takes hold of neuronal correlations in the network, leading to mean correlation *r* > 0.9.

Next, we examined whether the recurrent network simulation using the same network size (240 units) and a similar parameter setting (as presented in Fig. 2) could also fit the baseline shift and amplitude amplification of the slow spontaneous fluctuations observed during targeted free recall behavior, as reported in (Norman et al., 2017). In this regard, the parameters τ, *p* and *`μ`*_conn_ remained unchanged from the previous simulation.

Figure 3 compares the intra-cranially recorded HFB baseline shift amplification with the simulated neuronal activity. To that end, few remaining parameters of the simulated network were again fitted for optimal power spectrum match with results obtained by the intracranial HFB signal PSD. As for the setting of the decay constant, in addition to the optimal knee-location fit obtained in Fig. 2 (indicating τ = 20 ms), another approximation method was employed, a Lorentzian approach, following (Chaudhuri et al., 2018). The result of this approximation is shown in SI Fig. S3. It was established once again, now based on both methods, that setting this parameter near the biophysically-grounded-estimate of the membrane time constant optimizes the fit of the knee frequency location, and here also of the PSD curve for mid-range frequencies (> 0.1 Hz and < 10 Hz). To adjust the curve slope in the higher frequencies (> 10 Hz), the sampling fraction was set to α = 0.01. The simulation was run for 1800s, with the first 600s omitted as a stabilization period (by means of signal stationarity testing) before analysis of the remaining 1200s.

**Figure 3.**
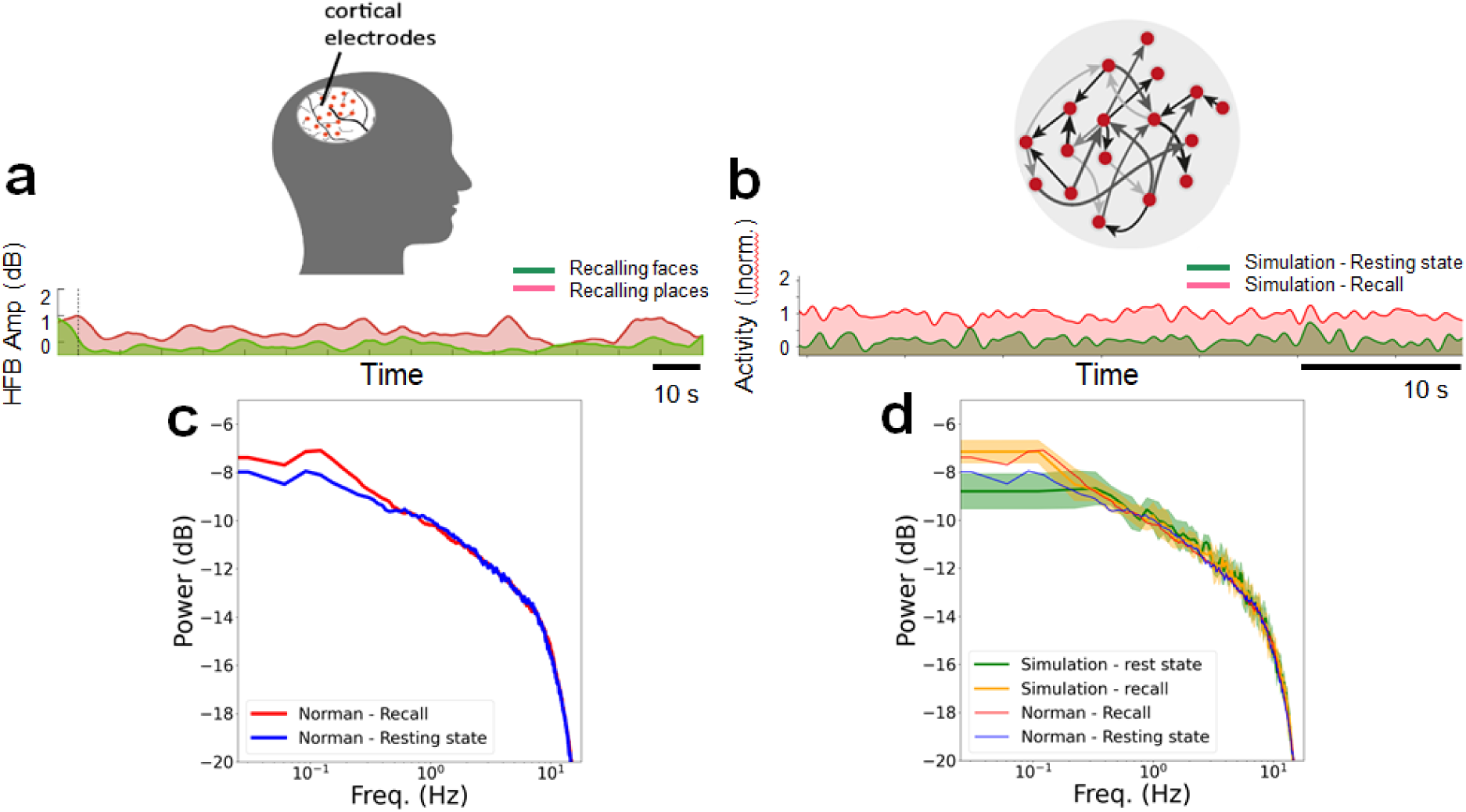
Amplification of slow-fluctuations in the brain and in the recurrent network. iEEG HFB signal from targeted free recall target (red) and non-targeted (green) neuron populations (adapted from Norman et al., 2017) in panel 3a, is compared with the simulated spontaneous activity in “recall” target (with additive noise) vs non-target “rest” blocks in panel 3b. The power spectra obtained from all intra-cranially recorded free-recall target sites (red) relative to the non-targeted population (blue) is shown in panel 3c. Simulated results are overlaid on the empirical PSD results in panel 3d, showing near fit, in expectation, over multiple realizations. The mean and standard deviation are shown for the free-recall target (yellow) and non-target (green).

Panel 3a depicts an example of the experimentally derived spontaneous HFB fluctuations in two neuronal populations - one that was targeted to elicit free recall (red) and the other that was not targeted (green). The baseline shift of the fluctuations for the targeted area is readily visible. It should be noted that in Norman et al. 2017, a generalized enhancement of slow fluctuations was found for the free recall conditions; however, the study did not find clear evidence for a selective enhancement in targeted vs non targeted categories, likely due to the mixed category selectivity and low SNR of the recording sites. By contrast, in the model simulation, we assume a fully selective neuronal group- and the simulation results should be taken as a prediction of such category-selective effects in future studies. For comparison, Panel 3b shows the simulated activity fluctuations for a network when a uniform baseline (fixed value) shift, *I*^*add*^(see Methods), was added to the white noise inputs (red) and without this added fixed value shift (green). Note that the gap between the two curves can easily be fine-tuned via setting the relative value and ratio of *I*^*ext*^ and *I*^*add*^, without compromising other measurable properties. Importantly, similar amplification in the baseline firing rate was observed also by increasing the network gain, *`γ`*, towards the critical point, implying a minor 1% change of *G*^∗^.

Panel 3c provides a more comprehensive analysis by showing the mean PSD across all intra-cranially recorded sites in the targeted neuronal population (red PSD) vs. the mean PSD across all sites in the non-targeted population (blue PSD). To make the comparison of these neuronal recordings to the simulations easier, we overlay the simulated results on top of the brain recordings in panel 3d. In this panel, the mean and standard deviation for multiple realizations (specifically, 8 arbitrary random-seed network connectivity initializations) of simulated target (yellow) vs non-target (green) blocks are presented. Here, the fit in PSD was reached by adding a fixed value shift *I*^*add*^of 12.5 pA to the white noise during simulated targeted (orange thick curve) but not to the non-targeted blocks (green thick curve). The control parameter was set within the *G*^∗^ range, as in Fig 2. As in (Norman et al., 2017), the increase in the power of the ultra-slow modulation in the simulated PSD amplitude during recall relative to rest blocks over the different realizations was statistically significant (for the range < 0.2 Hz; p < 0.05, FDR corrected, signed-rank test).

Importantly, a comparable amplification of the targeted ultra-slow frequency PSD was also achieved through direct modulation of the synaptic connectivity matrix (not shown). Notably, increasing the network gain, *`γ`*, by 1% within the scope of *G*_*R*_* was sufficient to replicate the empirically observed enhancement of slow frequencies during free recall. In SI Fig. S4 we show a distinct single realization PSD results from these simulations using the same color scheme mentioned above. Note the similarity in modulations of the PSD, and, specifically, the amplification in the ultra-slow-frequency domain of the PSD, evident in the targeted relative to non-targeted simulated blocks.

In the following subsections, we systematically examine the phenomenon of critical slowing down in our recurrent random network model, focusing on its fundamental properties rather than fitting specific data. Accordingly, some parameter settings are adjusted freely to enhance clarity in the illustrations.

To examine the impact of network connectivity on the PSD over a broader modeling range and greater detail we have systematically examined the influence of changing network and connectivity parameters over a wide range starting from well below the dynamical critical point. This is depicted in Fig. 4. In panels 4a-b, a simplified network setting with 100 units was used to explore the influence of the connectivity control parameter *G* on the spectra, via gradual wide range adjustments of *`γ`*, while keeping all other parameters fixed. The simulation lasted 800s, with the first 200s omitted in the analysis to allow signal stabilization; τ was set again to 20 ms, and α was set to 1.0.

**Figure 4.**
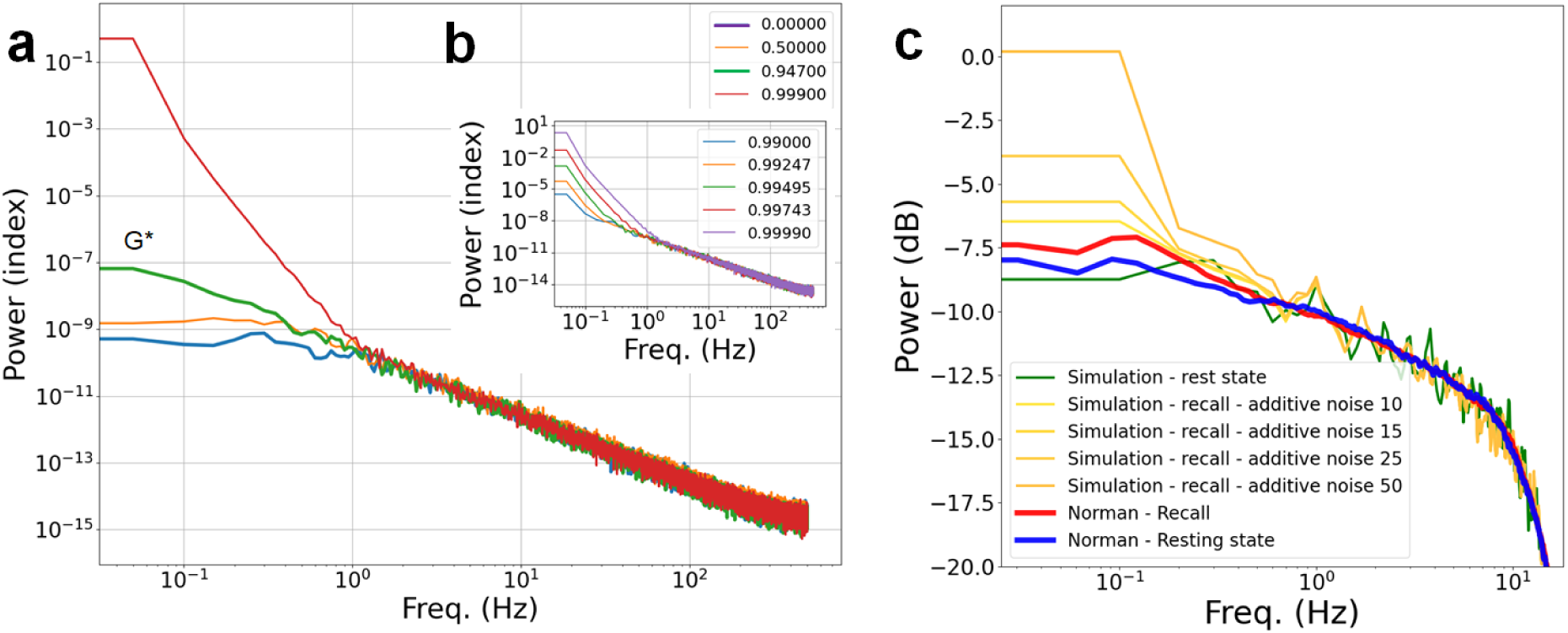
Effect of network connectivity strength and additive noise on the spontaneous fluctuations’ PSD. Panels 4a and 4b demonstrate the critical slowing down of low-range frequencies in an arbitrary network setting as the control parameter approaches the value of 1 (colors indicate different values of *G*). In panel 4a, the cases of 0-connectivity (i.e. γ = 0) and of *G*\* are included (blue and green bold, respectively). In the inset (panel 4b), the gradual verge-of-criticality slowing down is highlighted. Panel 4c shows a similar amplification of slow frequencies, here as a function of the amount of additive noise (see Methods *I*^*add*^) overlaid on PSD results from (Norman et al., 2017). Importantly, the network is set at the vicinity of criticality (*G*\*). Note that the different scales used on the ‘Power’ axis in the different panels are aligned with the relevant references (panels 4a and 4b are scaled to match Chaudhuri et al., 2018, whereas panel 4c follows the scaling used in Norman et al., 2017).

Panel 4a shows the effect of setting *G* over its full range from 0.0 to 1.0, whereas panel 4b focuses on the narrow range near the critical point. Three key findings emerge from this analysis: first, a gradual increase can be observed in the amplitude of slow frequencies that accelerates near the critical point of *G* (when *G* ≲ 1); second, the medium to high frequency range of the PSD remains unchanged even when all network connections are set to zero, effectively isolating the neurons from each other; and third, the slow-frequency PSD slope for *G*\* (green line), which is, interestingly, close to the scale-free exponent of negative 2. In panel 4c, the PSD results for the same simulation as in Fig. 3 are shown, with the control parameter in vicinity of *G*\*, but with gradually increasing levels of baseline shift of the noisy input, *I*^*ext*^. Similarly to the modulation of connectivity strength towards criticality in *G*, when the network is already near-critical, additive noise leads, as well, to a slowdown in the ultra-slow frequencies. To further investigate, we compared the influence of additive noise on the slowing down phenomenon in networks at varying proximities to the critical point. Figure SI S5 shows that additive noise significantly increases the amplitude of low frequencies when the network is near the critical point but has practically no effect when the network is far below the critical point.

The emergence of a scale-free power-law in the fluctuations of simulated activity (even in individual isolated neurons, for mid-to high frequencies) raises the concern that the network behavior merely reflected the dynamical structure of the injected input noise. To examine this possibility, we modified the injected noise profile by high-pass filtering and observed the effect on the network PSD for parameters near criticality. The simulation was 800s long, with the first 200s excluded for signal stabilization. The time constant τ was set here to 20 ms, as before. The control parameter, *G*, was set in vicinity to criticality and α was set to 1.0. Figure 5 displays the simulation results. We also used the same noise profiles to simulate isolated neurons (zero-connectivity network), with the resulting PSD shown in SI Fig. S6.

**Figure 5.**
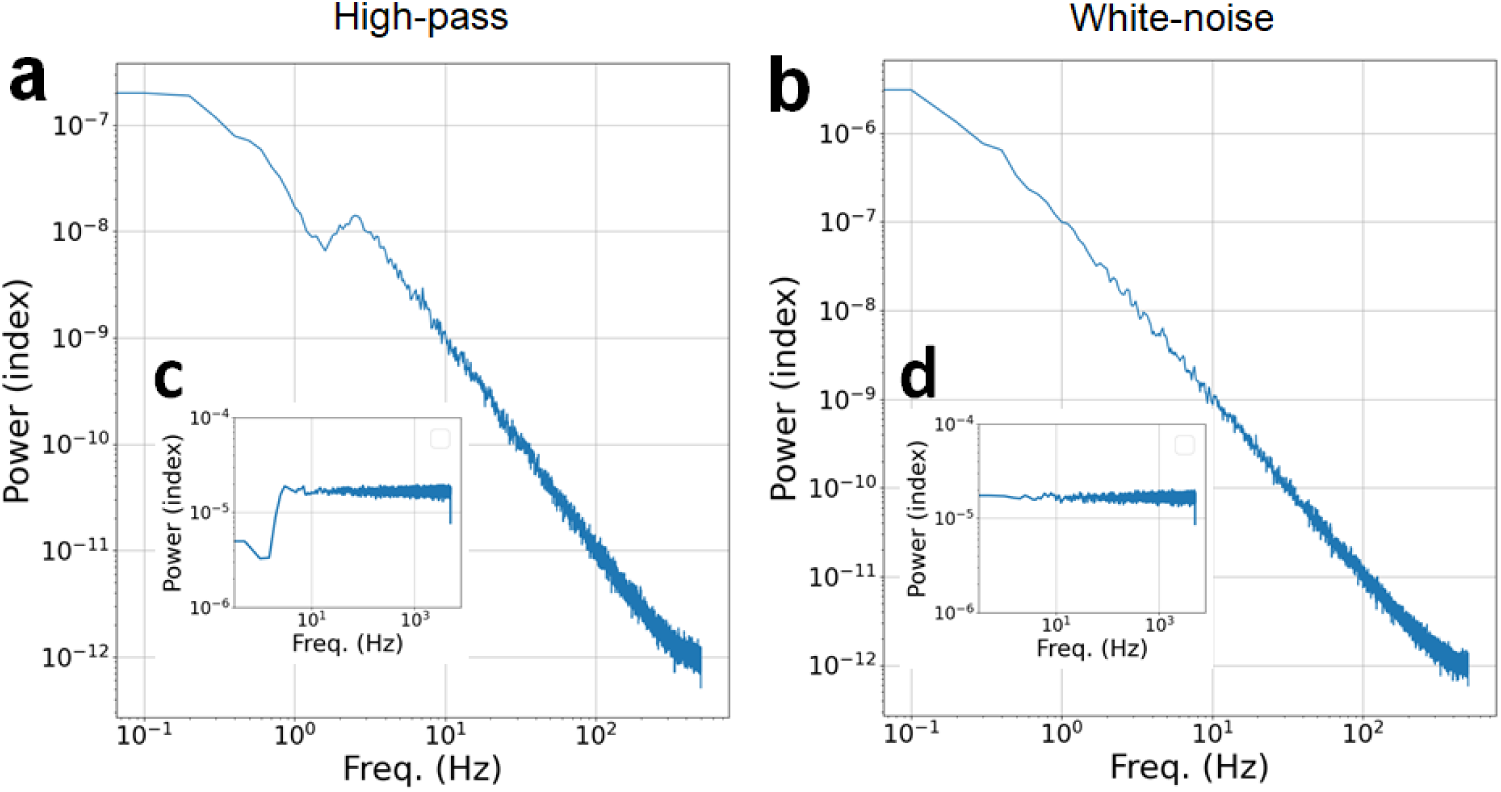
Impact of changes in input noise spectral profile on the network PSD. Panels a-b demonstrate the simulated network PSD when the injected noise is high-pass filtered (panel a) relative to the initial unfiltered white noise (panel b). Insets c – d present the noise spectra in use for each of the main panels, respectively.

As can be seen, even drastic changes in the spectral power profile of the input noise had only minor impact on the PSD mid-to high frequency power-law profile of the network as a whole. This is evident in the striking discrepancy between the PSD profile of the input noise (Fig. 5 insets c-d) and the PSD of the entire network (panels 5a-b). Of particular interest is the insensitivity of the network even when all slow input frequencies were drastically quenched to a tenth of their original power (see panel 5a relative to 5c). Similar behavior for the PSD of mid-to-high range frequencies is preserved also in the isolated neuron network (SI Fig. S5).

Finally, to analyze the influence of network size on the amplification of the spontaneous fluctuations’ slow frequencies as the network approaches criticality, we scanned the parameter space of both network connectivity and size. In the analysis, we iteratively generated network realizations of larger and larger scale (150 - 350 units), and increased the control parameter G (via *`γ`*) from subcritical to near-critical range, while collecting the slow (normalized by fast) frequency power values.

The results are shown as a two-dimensional heat map (Fig. 6). The x and y axes in this map depict network size and control parameter settings, respectively, whereas the color per pixel denotes the degree of slow frequency amplification (more details in Methods). The heatmap clearly demonstrates that modulation of the control parameter towards criticality is the dominant factor driving the slow-frequency power amplification effect. A mild trend for larger amplification in slow frequency power is shown for network size > 200. This trend appears to become negligible as networks grow larger.

**Figure 6.**
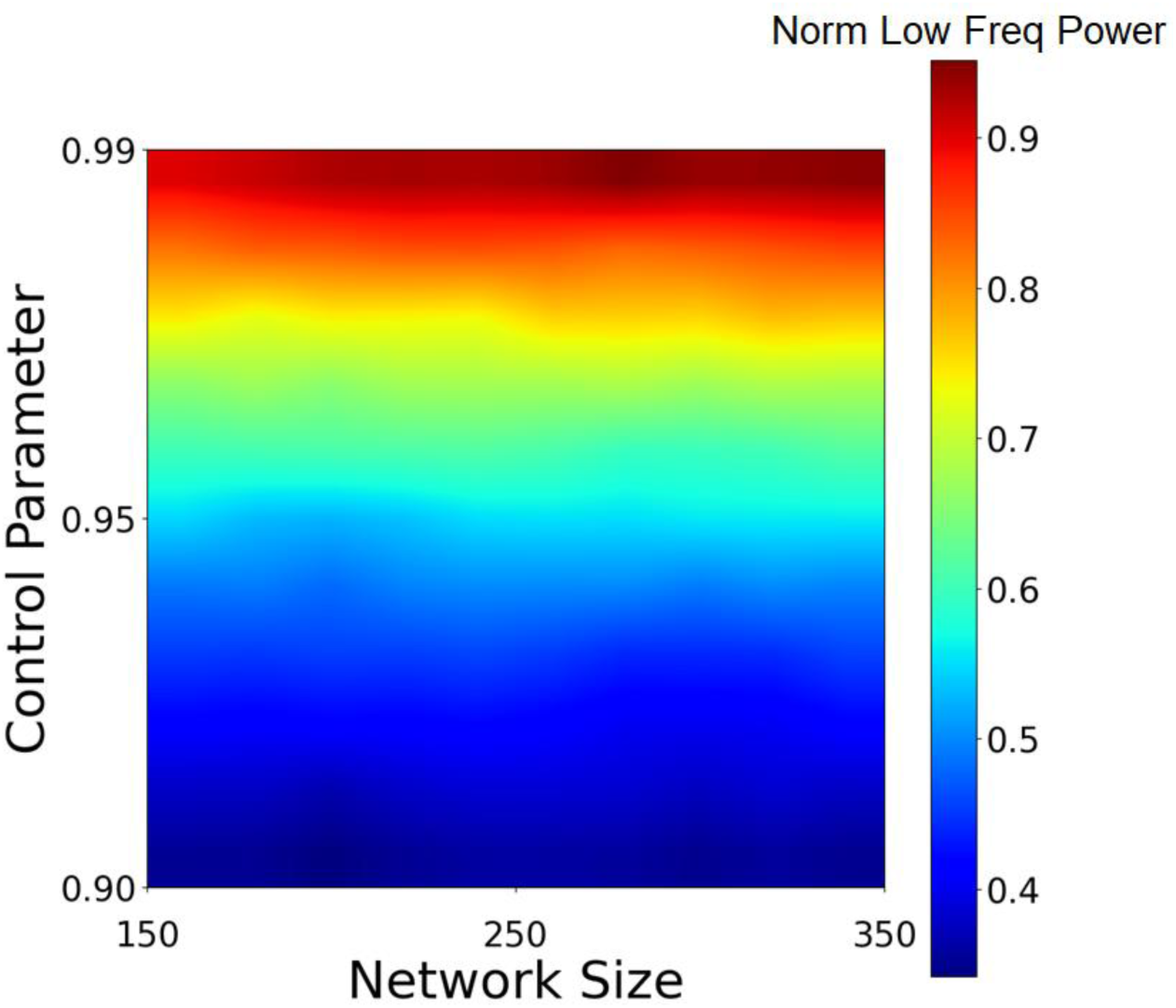
The impact of network size and network connectivity strength on the amplification of the slow spontaneous activity fluctuations. The map color scheme presents this amplification as slow frequency power, normalized by fast frequency power. Each of the cells represents an average of five random realizations, per specific network size and control parameter value.

## Discussion

Ultra-slow spontaneous fluctuations are one of the most widely studied phenomena of the human brain. Numerous studies have revealed that these fluctuations are organized along highly complex but informative functional connectivity patterns (Fox et al., 2006; Fox & Raichle, 2007; Sadaghiani & Kleinschmidt, 2013; Strappini et al., 2019) suggested to be related to cognitive habits, traits and pathologies (Eldar et al., 2013; Hahamy et al., 2021; Harmelech & Malach, 2013). It is also proposed that they drive self-generated, voluntary, and creative behaviors (Broday-Dvir & Malach, 2021; Malach, 2024; Norman et al., 2017; Norman & Malach, 2022).

Despite extensive research on the properties and dynamics of these slow fluctuations (Fox & Raichle, 2007; Nir et al., 2008; Raichle, 2011; Raut et al., 2021), the underlying neuronal mechanism remains unknown. The slow dynamics and intricate organization of these fluctuations may suggest a complex underlying mechanism, but we argue that the opposite is true. We propose that intrinsic fluctuations in recurrent networks exhibit a basic physical principle that leads to their slowing down and amplification of amplitude as the network approaches its critical point, a phenomenon known as critical slowing down. We hypothesize that this fundamental dynamics underlie the behaviors of the ultra-slow spontaneous fluctuations during both rest and task conditions.

Using a simple random recurrent network near criticality and unstructured white noise inputs, we successfully simulated the critical slowing down dynamics and reproduced the power spectrum of the spontaneous activity fluctuations as reflected in the intracranially measured HFB. Furthermore, a small uniform increase in synaptic efficacy, or alternatively, a small DC shift in this noise input, near the critical transition point of the network dynamics, were sufficient to simulate the amplitude enhancement documented during targeted free recall in the visual cortex. The idea that changes in the efficacy of recurrent interactions can modulate local intrinsic timescales was also explored in the context of selective attention (Zeraati et al., 2023). Specifically, their computational modeling work proposes that increased intrinsic timescales during selective attention are associated with a shift of network dynamics closer to a critical point. Recently, modeling of stochastic noise coupled with slow synapses in a recurrent spiking network successfully simulated the slow buildup anticipating voluntary movements (Gavenas et al., 2024).

It is important to clarify that our model was chosen not by virtue of its likely optimal fit to the experimental data - but rather because it was the most basic model of a recurrent network we could conceive of. The model was previously shown to account for the 1/f envelope of the power spectrum of the raw iEEG data (Chaudhuri et al., 2018). Furthermore, a major advantage of the simplicity of this model is that it allows for explicit control of the proximity to the critical point. By controlling the network gain, we can move a sub-critical model to a model that is near the critical threshold and explore the changes in the dynamics. In this respect, we examine a spectrum of models, namely various random recurrent networks, of different levels of connectivity and size that generate iEEG like activity, at varying proximity to the critical transition. The crucial point is that only under parameters that enable critical slowing down, namely near criticality, were we able to replicate the experimental data, and specifically the amplification in ultra-slow fluctuations.

Based on these simulations, we propose that the primary driver of slow spontaneous fluctuations in neuronal activity is stochastic, likely, but not necessarily, balanced noise, which then accumulates through the recurrent connectivity structure. The exact source of this noise is unknown and could originate from various processes. One possibility involves intrinsic neuronal mechanisms, such as thermal, synaptic, and ion channel fluctuations (Rusakov et al., 2020; Yarom & Hounsgaard, 2011). These processes could collectively contribute to the random membrane potential fluctuations consistently observed across all types of cortical neurons (Lampl et al., 1999; Raichle, 2011; Yarom & Hounsgaard, 2011). Another possibility is that the power spectrum of incoming spiking activity from subcortical sites exhibits characteristics of white noise. A recent study on activity in the subthalamic nucleus demonstrated that while the spectrum of synaptic activity follows a power law with an exponent around -2, the spectrum of the resulting spike rate is nearly flat (Liu et al., 2024). It is important to clarify, that our model equations capture the dynamics of synaptic activity, which is the primary contributor to iEEG signals (Einevoll et al., 2013).

In this work, we simulated two key features of spontaneous fluctuations: their power spectra and modulation during an intrinsically motivated task. However, we did not address how complex functional connectivity patterns emerge from such noise fluctuations. We have previously suggested that these patterns are shaped by synaptic structures and reflect prior learning (Harmelech et al., 2013; Harmelech & Malach, 2013). This hypothesis predicts that a recurrent network with updatable connectivity based on suitable learning rules will spontaneously reproduce a repeatedly activated specific pattern when network neurons, having learned the pattern, are injected with white noise near criticality. While we focus on the power spectra and modulation during task-induced targeting of categorical boundaries, future simulations could investigate the emergence of such functional connectivity patterns.

Our simulation focused specifically on ultra-slow fluctuations and their targeted amplification. While other aspects of the Norman et al., 2017 study are intriguing, they may require more sophisticated network models and additional analyses. The dynamics of the recurrent network when crossing the critical point, and its relationship with the concept of ‘ignition’ (Moutard et al., 2015), is another fascinating area of study. However, these aspects are beyond the scope of our current work and are left for future exploration.

As noted above, our model does not attempt to capture the full extent of the task and the consequent behavior. Rather, here we focus on the slowing down and amplification of the overall activity fluctuations. Our assumption is that shifting the network closer to criticality or enhancing the external noise may transition the network into a generative mode, where stored activity patterns are more likely to cross a relevant threshold. The advantage of the model in our context, is that the proximity to criticality can be directly controlled, allowing for a systematic examination of its effects on the resulting power spectrum.

Earlier studies have demonstrated the emergence of a scale-free 1/f power-law spectrum using a rate model simulation near criticality (Chaudhuri et al., 2018; Gao et al., 2020). However, they focused primarily on the tipping point of criticality (*G* ≈ 1) of the network parameter setting. In contrast, we aimed to investigate the minimal conditions required for the scale-free PSD to arise, examining network connectivity weights far below the critical point, as well as the impact of the input noise profile on the network’s neuronal activity.

Intracranial recordings of single neurons and the high frequency broadband (HFB) amplitudes of the LFP, which has been shown to index the average firing rate of cortical neurons (Manning et al., 2009; Nir et al., 2007), reveal the power spectra of the spontaneous fluctuations in the human cerebral cortex during rest (panels 2c and 3c). These spectra consist of two power-law exponents, with slower frequencies following a shallower exponent than higher frequencies. Remarkably, a similar profile emerges in a random recurrent network with fully stochastic connection strengths and input noise under the appropriate parameters (compare to panels 2d and 3d).

These results were obtained using minimal, biophysically-oriented, parameter settings, consisting of a physiologically plausible membrane time constant (∼20 ms), sampling of only a small part of the network nodes (α), which is compatible with the localized nature of the empirical recording sites, and control of the system’s slowing down by a single gain parameter (*G*).

We first discuss the high frequency slope of the HFB PSD. Analyzing the behavior of the network under zero connectivity (panel 4a) revealed that the resultant fluctuations in membrane potential approximated a power-law beyond a certain “knee” frequency, even when the spectra of the stochastic noise injected into each neuron was flat (white noise) or biased towards higher frequencies (see Methods and SI Fig. S5). Thus, each unit, when decoupled from the rest of the network, maintains the slow frequency band and attenuates the higher frequencies in a scale-free manner, following a Lorentzian power spectrum. A mathematical analysis presented in SI section 1 shows that the dynamics of a scale-free power spectrum with an exponent of negative 2 is indeed the expected result for the fast-mode HFB PSD. Recent studies of activity fluctuations in isolated cortical and hippocampal neurons also reveal that individual neurons are capable of generating stochastic fluctuations obeying scale-free power-law profiles as well as long time-scales, even in the absence of network connections (Gal et al., 2010; Gal & Marom, 2013a; Sukenik et al., 2021).

Examining the impact of network connectivity on the spontaneous fluctuations reveals a specific amplification of slow frequencies’ amplitude in the spontaneous fluctuations spectra as the network approaches its dynamical phase transition point. This is shown when separately examining the impact of changing the connectivity strength, i.e., the efficacy of intra-neuronal communication weight, and the richness of connectivity as reflected in network size (Fig. 6). A successful amplification of slow frequencies can only be achieved within a regime in which these two parameter spaces overlap. However, only random network configurations were explored in this study, and a better structured connectome arrangement, whether shaped by innate prearrangement or as learned by prior experiences, may present other near-critical behaviors. Nevertheless, it should be noted that even in extremely small, properly configured networks possessing fewer than 20 neurons, a powerful amplification of ultra-slow fluctuations can be obtained, suggesting that ultra-slow fluctuations lasting many seconds do not necessarily depend on large-scale cortical networks for their generation, as also suggested by previous work (Gal & Marom, 2013b).

Our simulation results are compatible with the notion that the spontaneous fluctuations can be selectively amplified when behavior switches into a free recall state. Furthermore, they predict that when participants attempt to visualize different items belonging to a certain category of visual stimuli, their slow spontaneous fluctuations should be selectively amplified, specifically in the corresponding neuronal representations. This can happen through a modulation towards criticality via connectivity gain or via additive external noise (see panels 3a and 3c and more details in Norman et al., 2017). Such amplification enhances the probability of these fluctuations to cross the decision bound in the category-selective area. This modulation toward a critical threshold provides an elegant mechanism for rapid and flexible control over the expression of any spontaneous behavior within an intentionally selected categorical boundary. For example, when we attempt to freely imagine different types of faces, we rarely, if ever, mistakenly picture a building. As we demonstrate here, modulation of proximity to criticality and the resulting critical slowing down offer a plausible mechanism for such precise cognitive distinctions. However, it should be emphasized that such specific, category selective, enhancement of slow frequencies has not been demonstrated experimentally; yet, likely due to low selectivity and poor SNR or the recordings available in Norman et al’s 2017 study. Thus, our simulation of this result should be taken as a prediction for future studies with a higher spatial resolution.

When local populations’ activity fluctuations transiently cross the decision bound, this may lead to rapid self-amplification termed “local ignition”. Such ignitions have been experimentally observed during crossing the perceptual awareness threshold using iEEG recordings (Fisch et al., 2009; Moutard et al., 2015). Additionally, near a critical point, network sensitivity increases, enabling better encoding and recall of information through balanced response to inputs and adaptation to noise (Kim et al., 2021; Shriki & Yellin, 2016).

Our simulations of a random recurrent network demonstrate that a simple shift in noise level or in gain modulation accurately reproduces the intra-cranially recorded slow frequency amplification observed in the empirical data during category-selective visual recall (panel 3d and SI Fig. S4, respectively). The key factor inducing the amplification effect is the network’s proximity to its critical transition point, regardless of the specific mechanism controlling its excitation.

It is important to note that empirical brain data and recurrent network simulations specifically relate to fluctuations in neuronal firing rate or, in cases where such measures are unavailable, to the broadband high frequencies (gamma) envelope of the LFP. Previous studies have shown that the gamma envelope reflects fluctuations in the average firing rates of local neuronal populations (Manning et al., 2009; Nir et al., 2007). These measures of firing rate fluctuations should be distinguished from fluctuations in the local field potential, i.e., the LFP, itself. Thus, it should be emphasized that the power-spectrum of the LFP should not be confused with the power spectrum of the firing rate or HFB fluctuations, as the two types of fluctuations can be antagonistic to each other under different experimental paradigms (Privman et al., 2011).

Moreover, previous studies have suggested that the LFP power spectrum is optimally fitted by setting the dynamics control parameter precisely at the tipping point of criticality (i.e. at *G* = 1), which could potentially imply fundamental instability and a network easily crossing between subcritical to supercritical states (Chaudhuri et al., 2018; Gao et al., 2020). However, our optimal fit for the HFB power spectra implies that relevant cortical networks operate below this critical transition point (i.e., at *G*^∗^∼0.94), where stability is still maintained but close enough to critical dynamics to allow the system to computationally benefit from near-criticality effects.

Wider implications of the critical slowing down phenomenon should be noted. For example, it is well known that sleep deprivation can lead to over excitability and even to epileptic seizures. Indeed, it is proposed that sleep deprivation influences the proximity to criticality (Harel et al., 2025; Meisel et al., 2013). It is also known that sleep deprivation slows down reaction times and increases the presence of slow frequencies (Harel et al., 2025; Meisel et al., 2015; Nir et al., 2017). These phenomena may all be related. It may be that extensive recall tasks, which bring the system closer to criticality, will not only manifest in critical slowing down but even trigger seizures in those who are susceptible.

A key hypothesis of our study is that cortical activity is modulated closer to a critical point in order to shift to a more generative mode, which is required under a wide range of conditions, such as creative and associative thinking (Malach, 2024; Norman et al., 2017; Pezzulo et al., 2021). The prediction is that behaviors requiring a generative mode would lead to slowing down of the dynamics compared to resting state. Note that this may potentially manifest in other signatures of criticality, such as measures related to edge-of-chaoes dynamics and neuronal avalanches.

Finally, it is worth noting that the relationship between the LFP and firing rates is more appropriately captured by measuring the changes in the exponent of the LFP power-law PSD, as observed in our previous intra-cranial analysis of visual responses (Podvalny et al., 2015). While these issues could potentially be examined in our recurrent network simulations, they have not been pursued in this work.

Overall, the findings from our simulations suggest that the ultra-slow spontaneous fluctuations are an emergent network phenomenon and support the idea that the cortex operates near a critical point, allowing for rapid and flexible control over spontaneous behaviors while maintaining stability. Furthermore, they provide a mechanistic explanation for the ability to flexibly target specific categories for free behavior. This work makes specific predictions and opens new avenues for investigating the role of criticality in brain function and could potentially lead to the development of new computational models for understanding the neural basis of cognition and behavior.

## Methods

### Random recurrent network as a rate model

Rate models can be derived from the well-studied Wilson-Cowan equations (Wilson & Cowan, 1972). They were explored within the context of iEEG data in recent literature (Chaudhuri et al., 2018; Gao et al., 2020). In previous work (Shriki et al., 2003), we have shown that such rate models, describing the neuronal firing rates, can be faithfully derived from large-scale conductance based networks resembling cortical organization. The simplicity of rate models makes them highly useful in studying properties of large networks.

Here, we considered a random recurrent network model consisting of N rate-based units representing single neurons or groups of neurons (see Figures 2-6). We closely follow the model proposed in (Chaudhuri et al., 2018) to explain the power spectrum of iEEG recordings. The rate dynamics of the *j*-th unit over time in this model is given by:

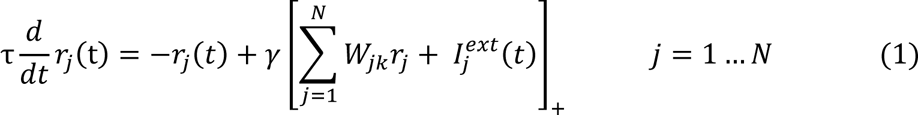

where *r_j_* is the *j*-th unit activity level, and *τ* is a characteristic integration time. The [ ]_+_ brackets denote a rectified linear (ReLU) activation function. The γ variable denotes the slope of the firing rate – current curve (f-I curve). The weights *W*_*jk*_ describe the strength of the connection from the presynaptic unit k to the postsynaptic unit *j*, and 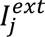 denotes the external input to the *j-th* unit (drawn here randomly from a “standard normal” distribution).

The dynamical variables, *r_j_*, can be interpreted as representing synaptic variables, which effectively low-pass filter the neuronal firing rates (Shriki et al., 2003).

### Criticality in the random recurrent network

We consider a network architecture comprising a non-hierarchical cluster of recurrently connected neurons characterized by a semi-linear f-I curve with gain γ. Following (Chaudhuri et al., 2018), we choose the connections in this network (the entries of matrix *W*) to be sparse and random. More specifically, each entry is nonzero with probability *p*, and the nonzero entries are drawn from a normal distribution, with positive mean μ_conn_ and variance σ_conn_, normalized by the size of the network, N.

When set as a rate model (assumed to be represented by the iEEG high-frequency broadband power modulation), we hypothesize it depicts the summed recurrent synaptic current into a fraction of neurons in this network, along with an intrinsic complementary membrane noise or some form of external noise.

Previous work (Chaudhuri et al., 2018) demonstrated how the product of three main parameters controls the proximity to the network’s critical point. These are: μ_conn_, γ, and *p.* At the tipping point of criticality, they satisfy the relation γ*p*μ_conn_ = 1, see (Chaudhuri et al., 2018). Thus, we define a control parameter G = γ*p*μ_conn_, and require that it lies in the vicinity of the critical point - G ≲ 1. In a near-critical state, either one of these arguments composing the control parameter could be varied to yield similar results towards the crossing point into a supercritical state.

To provide additional insight in the present context, we note that near a steady-state, an algebraic linearized approximation of the spontaneous fluctuations dynamics can be obtained. We briefly summarize the details given in (Chaudhuri et al., 2018; Ganguli et al., 2008):

1. The eigenvalue spectrum of such random matrices is composed of a cluster of complex eigenvalues with negative real part, and a single isolated eigenvalue, which corresponds to the uniform eigenvector (namely, to the mean network activity).
2. This matrix that controls the linearized dynamics is a random matrix whose dominant eigenvalue, *λ*_*dominant*_ = −1 + *G*, converges ever closer to 0 as the system approaches its bifurcation point at *G* = 1. At this point, the system transitions from stable to unstable dynamics.

Importantly, the underlying analysis relies on random matrix theory and consequently assumes that the network has a large scale (Rajan & Abbott, 2006; Tao, 2013), specifically, a minimum of a few hundred units (see SI Fig. S1). This scale is fundamental because random matrix theory draws on statistical properties that accurately reflect the system’s behavior only when applied to large matrices. Adequate network size is thus vital for capturing the delicate dynamics and stability features essential for close analysis of proximity to criticality.

Critical slowing down is manifested in the system’s temporal dynamics, as they are linked to the inverse of the abovementioned dominant eigenvalue (see SI for additional detail). As the system approaches criticality, signals reverberate in the network for increasingly longer time, producing a slowly decaying autocorrelation function and increasing the power of low frequencies.

For the sake of convenience, the following parameters were kept fixed across all simulation runs presented in the paper: μ_conn_ = 49.881 pA/Hz, σ_conn_ = 4.988 pA/Hz, *p* = 0.2. Unless otherwise noted, the parameter that was used to control the proximity to criticality was the slope of the f-I curve, *γ* (for *γ* ≈ 0.1, we obtain *G* = γ*p*μ_conn_ ≈ 1). We did however verify that qualitatively equivalent results can be reached by modifying G using other parameters.

### The characteristic integration time and PSD profile’s pattern

As for the characteristic integration time τ, we explored values in the range of 2 – 200 ms. Using a simple linearized mathematical approximation (see SI Fig. S3 and (Chaudhuri et al., 2018)) it can be shown that the commonly observed “knee”-bend in in the power spectral density (PSD) profile appears near 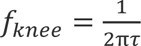, and that modulation of slow-frequencies is also significantly influenced by this the integration time constant, implying that the fit tuning process should carefully consider its setting.

Our findings here indicate that using the previously reported timescale of the membrane time constant in the proximity of τ = 20 ms (Gao et al., 2020; Koch et al., 1996; Softky & Koch, 1993) led to an optimal fit regarding the location of this knee in the activity rate PSD curve as well as the related slow frequencies slope. That is, for both of the cases explored in (Nir et al., 2008; Norman et al., 2017), as demonstrated in Fig. 2 and 3 respectively.

The parameter α, defined as the fraction (i.e. 0 < α < 1) of network nodes selected for summation, plays a significant role in determining the slope of medium range frequencies (> 0.1 Hz and < 10 Hz) in the PSD curve, since it influences the mixture of slow and fast dynamical modes exposed in the system. When constructing the PSD, we sum the activities of α*N* random nodes and compute the spectrum of the summed activity. The larger the value of α, the larger the contribution of the dominant slow mode to the mid-range PSD slope, since this mode is shared among the network nodes.

### Simulation and parameter fitting efforts

Simulation code, including full implementation of the rate model and plotting of figures, was developed in Python, taking advantage of suitable open-sourced libraries’ methods (including, but not limited to, SciPy and Matplotlib), facilitating tuning efforts for fitting the empirical results (details below).

To determine a stationary signal (and the period to be omitted before signal stabilization), the ‘Augmented Dickey-Fuller’ method was iteratively applied, using the Python statsmodel library’s adfuller method, extending the delay, until the remaining segment’s stability reached p-value < 0.01. For the results depicted in Fig. 2, the signal was low-pass filtered (for < 0.1 Hz) as in (Nir et al., 2008), using a Butterworth method. Correlation, auto- and cross-correlation coefficients were computed using time-lagged correlation methods. In panel 2d, the activity of two arbitrarily selected highly-correlated neurons is presented. In panel 2e, the power spectrum was computed using Matplotlib mlab’s PSD, implementing the Welch method.

A simple amplitude scaling factor k was applied to match the Welch method results with those presented by direct use of FFT power in (Nir et al., 2008). A regression line curve was fitted using a curve-fitting method in the range of 0.1 – 10 Hz. Following Nir et al., 2008, the autocorrelation was computed using signals from the first sampling only (resembling a single cluster of neurons in the network), and cross-correlation was computed iteratively between neuron pairs of both samples. Finally, the power spectrum was obtained by computing the mean of these autocorrelation (black line) and cross-correlation (orange line) curves, plotted in log scale.

For Fig. 3, the simulation was arranged as consecutive “rest” vs “free-recall” 1800 s blocks. Additive noise *I*^*add*^ was defined as constant value within each simulation run, so that 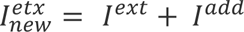. For the purpose of sampling a satisfactory distribution for statistical analysis, the simulation was repeated using different random seed values. The multiple realizations generated were independently compared with the empirical data. Other than that, similar simulation and computational methods were applied as in Fig. 2. Testing for various possible confounds, the simulation configuration for the results presented in Figures 4 – 6 was suitably adapted, but remained similar in concept to the settings in Fig. 3. In data preparation for Fig. 5, white noise was sampled in advance, applying suitable filters (using scipy signal.butter methods) over the random (white noise) normal distribution based temporal signal. For the map presented in Fig. 6, we iterated through the parameter space of network connectivity and size, iteratively generating random network realizations of larger and larger scale. Thus, as the control parameter G was modified (via *`γ`*) from subcritical to near-critical range, values of the slow (normalized by fast) frequency power modulation were collected.

### The human iEEG data and its analysis

The empirical data used in this study has been previously reported in (Nir et al., 2008 and Norman et al., 2017). In both studies, intracranial recordings were obtained from patients with pharmacologically resistant epilepsy who were undergoing a surgical evaluation process for the treatment of their condition. In the first study (Nir et al., 2008), data were recorded from five participants (over multiple sessions) under the conditions of wakeful rest. During the wakeful rest, subjects’ eyes were closed and the room was dark and quiet, however, alertness was occasionally monitored. The second study (Norman et al., 2017) included twelve participants who performed a free recall task, as follows. Immediately after a 200 s period of closed-eyes resting state, participants were presented with 14 different pictures of famous faces and popular landmarks in a pseudorandom order (each picture was repeated four times, and shown for 1500 ms, with 750 ms inter-stimulus interval). After viewing the pictures, participants put on a blindfold (or closed their eyes) and began a short interference task of counting back from 150 in steps of 5 for 40 s. Upon completion, recall instructions were presented, guiding the patients to recall items from only one category at a time, starting with faces in one run and with places in a second run. Participants were asked to verbally describe each picture they recall, as soon as it comes to mind, with 2–3 prominent visual features. The duration of this free recall phase was 2.5 min per each category. In both experiments analyses of iEEG data were focused on changes in high-frequency broadband amplitude (HFB, also known as high-gamma, defined as the range 40-100 Hz in the first study and 60-160 Hz in the latter one), shown to be a reliable marker of local neuronal population activity (Nir et al., 2007).

## Notes

### Competing Interest Statement

The authors have declared no competing interest.

### Summary of Updates

This version of the manuscript has been revised to update the Abstract and Introduction, incorporating new figures in the main text and SI, and enhancing our manuscript with additional statistical analysis.

