## Supplementary Information for "Adaptive proximity to criticality underlies amplification of ultra-slow fluctuations during free recall"

**Supplementary Information for “Modulation of proximity to criticality enhances slow activity fluctuations during free recall”** (Yellin, Siegel, Malach and Shriki)

1. **The power spectrum of a single (decoupled) neuron at high-frequencies**

The rate dynamics of single unit without any interactions are described by:

$$\tau\frac{dr}{dt}=-r+\gamma I^{ext}$$

Applying Fourier transform to both sides of the equation, we obtain:

$i\omega\tau\hat{r}=\hat{r}+\gamma\hat{I}^{ext}$,

where we have used the fact that the transform of the derivative is $i\omega$ times the transform of the function. This gives a closed form expression for the power spectrum:

$$\hat{r}(\omega)=\frac{1}{1+i\omega\tau}\gamma\hat{I}^{ext}(\omega)$$

Namely, the spectrum of the input is multiplied by:

$$Q\left( \omega\right)=\frac{1}{1+i\omega\tau}=\frac{1}{1+\omega^{2}\tau^{2}}\sqrt{1+\omega^{2}\tau^{2}}e^{i\phi}=\frac{e^{i\phi}}{\sqrt{1+\omega^{2}\tau^{2}}}$$

Where - $\phi=\tan^{-1} \left( \omega\tau\right)$

The corresponding power spectrum is:

$$\left| Q\left( \omega\right) \right|^{2}=\frac{1}{1+\omega^{2}\tau^{2}}=\frac{1}{1+{4\pi^{2}f}^{2}\tau^{2}}\sim\underset{\text{at high frequencies}}{\underbrace{\frac{1}{f^{2}}}}$$

Thus, for a white noise input (uniform spectrum) we obtain the spectrum above, which is flat at low frequencies and has a power law profile with an exponent of negative 2 at high frequencies.

2. **The effective proximity-to-criticality control parameter as a function of network size**

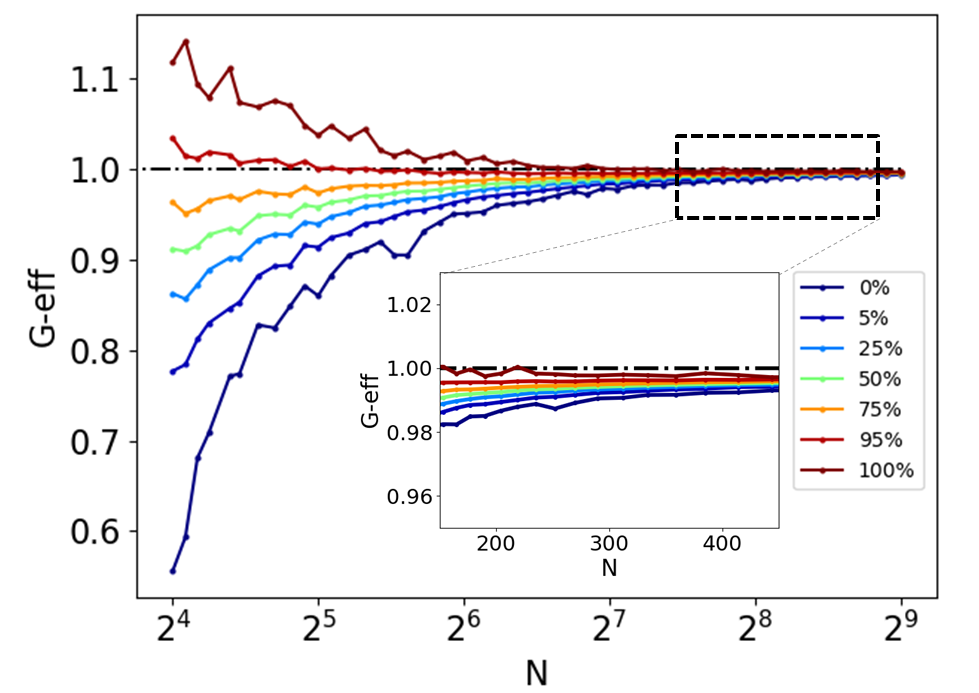

**Figure S1. Relationship between network size and the effective control parameter (**$\mathbf{G}_{\mathbf{eff}}$**), illustrating the variability in proximity to criticality for small networks.** The real part of the largest eigenvalue in the linearized matrix representation determines the network gain. To achieve critical dynamics, the effective gain should be 1. The figure shows the distribution of the effective gain as a function of network size when the parameters are set to yield a theoretical value of 1. Quantiles represent 100 random realizations of the network connectivity for each distinct network size. The analysis clearly demonstrates how the variance diminishes with increasing network size. Note (in the inset) that when the network size exceeds 200 units, the effective gain is very close to 1. For small networks, the network can be effectively sub- or super-critical, even when theoretically set to be near-critical. These results stem from the fact that the relevant analytical findings regarding random matrices hold only in the limit of very large networks.

3. **Exploration of various settings for the membrane time constant**

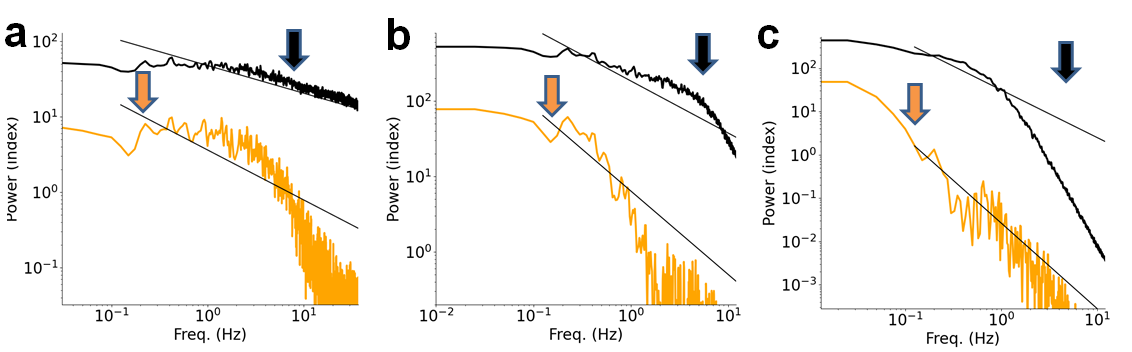

**Figure S2. Different settings of** τ **in fitting the PSD curve knee frequency.** Parameter fitting of Nir et al., 2008 empirical iEEG HFB signal results, using an artificial neural network model with 400 neurons, explored the optimal setting for τ. The autocorrelation (black line) and cross-correlation (orange line) PSD curve data, from two highly correlated neurons in the artificial network, are presented in panels a - c for different values of τ, 2, 20 and 200 ms, respectively. As indicated by arrows pointing to the knee position within the empirical autocorrelation (black) and cross-correlation (orange) PSD curve data, only panel b (reflecting τ =20 ms) demonstrates an accurate match.

4. **A linear system approximation of the empirical PSD results**

The Lorentzian approximation guides us towards rough estimates for the $\alpha$, *f*_slow_ and *f*_fast_ parameters. Here, *f*_fast_ is chosen to best fit the HFB signal knee frequency, in accordance to $f_{knee}=\frac{1}{2\pi\tau} \approx8 Hz$ (using $\tau=20 ms$), and α is chosen to match a suitable mix of the 2 terms (for fast and slow modes) – here set to 0.0025, in the Lorentzian fit.

$$\mathrm{PSD}\left( f \right)= K\left( \frac{\alpha}{f^{2}+ f_{slow}^{2}}+ \frac{1- \alpha}{f^{2}+ f_{fast}^{2}} \right)$$

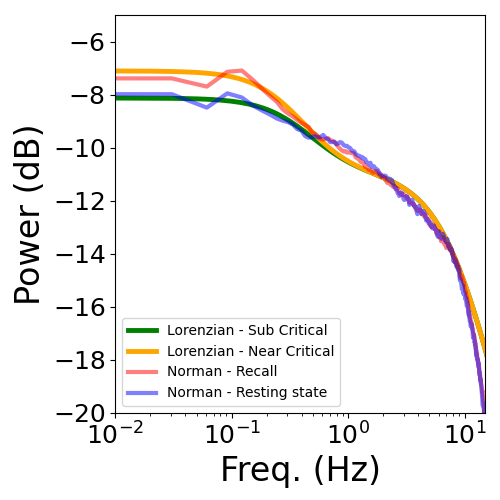

**Figure S3. A Lorentzian fitting of free recall vs. rest HFB PSD curves.** The power spectrum of the Lorentzian function is near-flat at low-frequencies and shows $\frac{1}{f^{2}}$ scaling at high frequencies, with a transition point set by the time-constant of the exponential. The fitting process maximized correlation between respective Lorentzian and HFB PSD curves. The different fitting for the sub- and near-critical curve fits lies in the setting of *f*_slow_ which was slightly adapted.

5. **Amplification in ultra-slow fluctuation power due to additive noise and gain control**

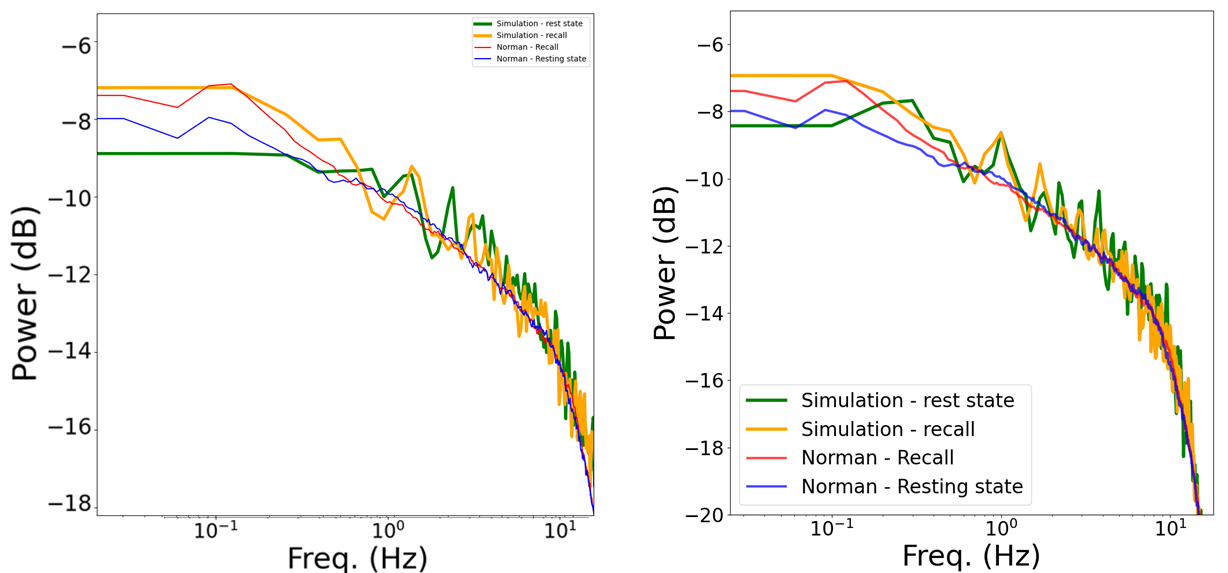

**Figure S4. Slow-fluctuations power increase in iEEG and in the recurrent network simulation.** Single realization simulation results from the fitting of Norman et al 2018 free-recall vs. rest PSD curves are demonstrated using additive noise and gain control**.** In both panels, the iEEG power spectra obtained from empirically recorded free-recall target sites (red), relative to the non-targeted population (blue), are compared with respective simulated results. In the left panel, additive noise is used to modulate slow fluctuation power increase whereas the F-I curve gain parameter, γ, is modulated for this purpose in the right panel. Note that Panel 4c in the main text extends the fit shown here by demonstrating the influence of higher levels of additive noise on the amplification in power of the slow fluctuations.

6. **The impact of additive noise**

Significant amplification of ultra-slow frequency power occurs only when additive noise increases in the immediate vicinity of the critical transition point**.**

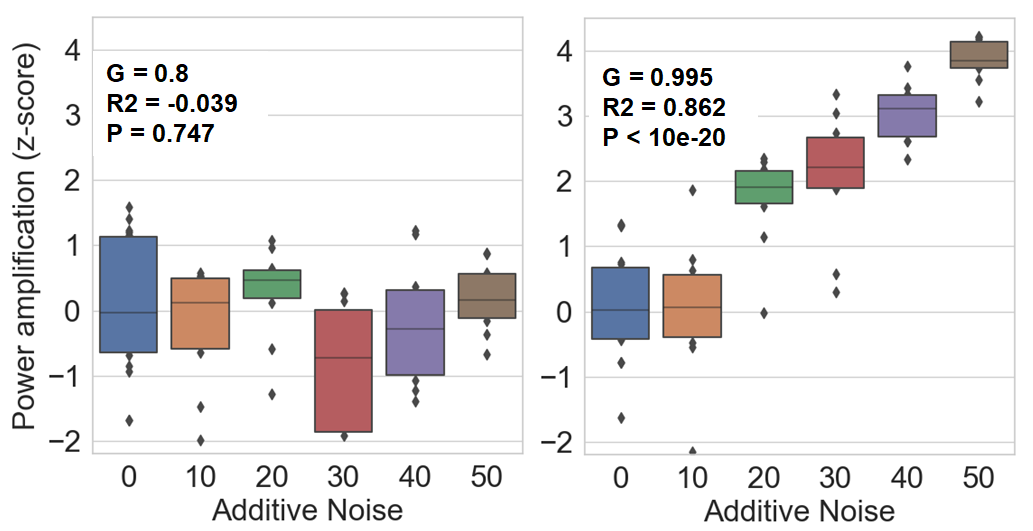

**Figure S5.** **The influence of additive noise on the ultra-slow frequency power.** Simulation results from our recurrent random network model far below the critical point (left panel) and at its very near vicinity (right panel) indicate that the significant amplification of the ultra-slow frequency power due to increase in additive noise is dependent on the proximity to criticality. In both panels, the x-axis indicates additive noise and y-axis denotes the normalized slow-frequency power amplification. Results are based on 10 random realizations for each level of additive noise. Normalization is two-staged, first obtaining the ratio of slow over fast frequency power per each realization, and then dividing the obtained value by the 0-additive-noise mean. Statistical analysis indicates a significant effect only for the case of near critical simulation results (R^2^ = 0.862, p < 10^-20^).

7. **Scale-free spectral profile is preserved when high-pass filtering the input noise in a zero-connectivity network**

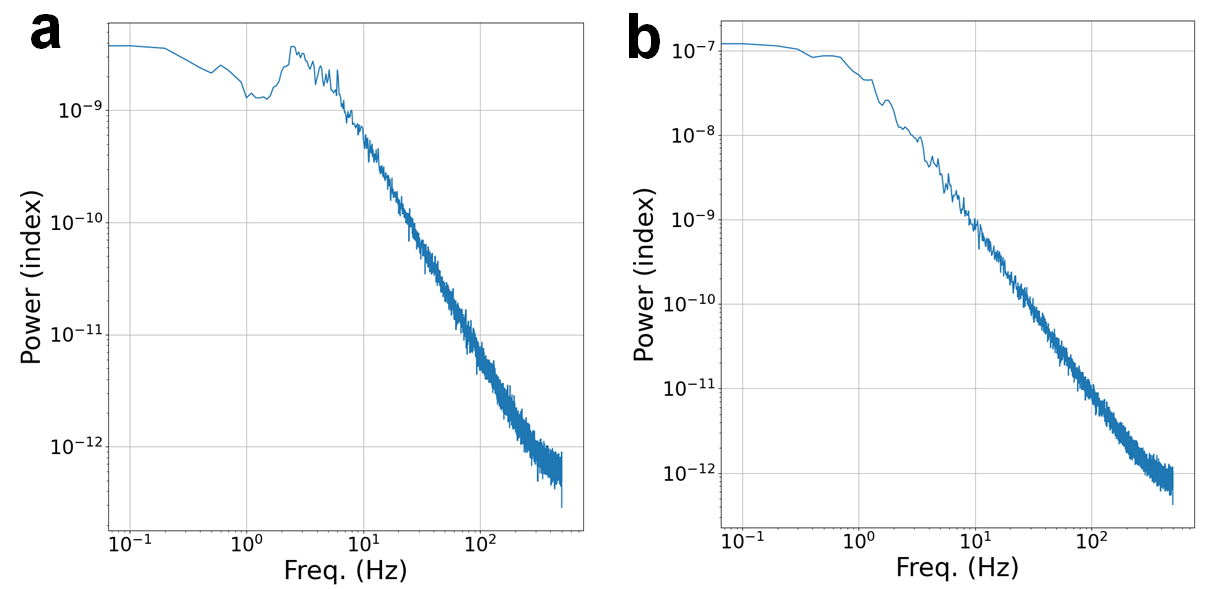

**Figure S6.** **Impact of different input noise spectral profiles on the PSD of a zero-connectivity network.** Panels a-b demonstrate a simulated zero-connectivity network PSD when the injected noise is high-pass filtered (panel a) relative to the initial unfiltered white noise (panel b). The cutoff frequency was 2 Hz for the high-pass filter. As in Figure 5 of the main text, the scale-free spectral profile in the mid-to-high frequency range is qualitatively preserved.
